# CAGE-TRX expands the scope of time-resolved crystallography through genetically encoded active-site photocaging

**DOI:** 10.64898/2026.06.26.734858

**Authors:** Clyde A. Smith, Ailiena O. Maggiolo, Chasity Janosko, Maura E. Charette, Nilakshi Paul, Marta Toth, Guillermo Calero, Sierra M. Carr, Silvia Russi, Sergei B. Vakulenko, Alexander Deiters, Aina E. Cohen

## Abstract

We present CAGE-TRX, a broadly applicable time-resolved strategy for *release-quench-probe* cryocrystallography and *release-probe* room temperature serial crystallography. These workflows enable light-triggered control of enzyme activity via genetically encoded photocaged amino acids. By decoupling reaction initiation from substrate design, this approach allows synchronized catalysis *in crystallo* and the capture of transient intermediates. Using β-lactamases as model systems, we demonstrate efficient decaging, restoration of activity, and structural visualization of reaction intermediates.

## Main

Enzyme catalysis is inherently dynamic, spanning conformational changes across multiple length and time scales, from subtle reorientation of active-site residues to loop motions that assemble or occlude catalytic pockets, and larger domain rearrangements and partner interactions. These motions are essential for function, linking structure directly to catalytic mechanism, evolutionary adaptation, and regulatory control. Capturing such dynamics remains challenging. Conventional cryocrystallography^1,2^ has provided atomic-resolution structures of countless macromolecules but largely yields static snapshots that obscure the conformational heterogeneity and transient states central to catalysis. Flash-cooling arrests motion, often fixing residues in single conformations that may not fully represent their functional roles. Although room-temperature (RT) crystallography can reveal limited ensemble behavior^3^, time-resolved crystallography (TRX) enables direct visualization of enzymatic processes in four dimensions^4-11^, providing unique insight into catalysis in motion.

All TRX experiments require a well-defined starting state and a means of initiating the reaction *in crystallo*, followed by structural interrogation at discrete time points along the reaction trajectory. Existing triggering strategies include rapid mixing approaches^12-14^, mix-and-quench methods^15-17^, and light-based initiation using intrinsically photosensitive proteins or photocaged substrates^18-20^. Among these, photocaging offers particularly precise temporal control, especially when the caged ligand is pre-positioned in the active site. A diverse set of caged molecules have been developed to target functional groups such as phosphates, carboxylates, thiolates, and phenolates, with caged nucleotides (e.g., ATP and GTP) among the most widely used (Supplementary Figs. S1a, S1b). The photochemistry underlying their activation is well established^18,21^. However, despite these advances, TRX applications remain somewhat limited by the narrow range of systems compatible with caged substrates, the difficulty of achieving uniform reaction initiation in crystals, and the synthetic complexity required to generate suitably caged compounds. Consequently, only a small subset of enzymatic reactions can currently be interrogated using light-triggered approaches. A broadly applicable strategy that bypasses substrate modification would substantially expand the range of systems accessible to TRX analysis.

To address this limitation, we introduce CAGE-TRX (Caged Amino acids for General Expansion of Time-Resolved Crystallography), a strategy that uses genetically encoded photocaged amino acids to directly control enzyme activity in protein crystals. In this approach, an essential active-site residue is replaced with a photocaged analogue via unnatural amino acid (UAA) mutagenesis^22-25^ (Fig. 1a), rendering the enzyme inactive until light-induced decaging initiates catalysis (Fig. 1b). Current UAA mutagenesis methodologies allow photocaging of several residue classes, including lysine, tyrosine, cysteine, serine, histidine, phenylalanine, glutamate, and aspartate^25^, extending the approach to diverse catalytic mechanisms and active-site architectures. In the CAGE-TRX approach, crystals of the caged enzyme can be prepared in the presence of substrate, cofactor or inhibitor, and a UV flash defines the reaction start point. This strategy is compatible with multiple time-resolved crystallographic workflows, including *release-probe* experiments (such as on-the-fly serial measurements at room temperature), and, as described here, a *release-quench-probe* approach, where substrates, cofactors or inhibitor are introduced into preformed inactive substrate-free or apo crystals at room temperature, and then freeze-quenching the crystals at different timepoints after reaction initiation to enable the capture of structural intermediates along the reaction coordinate.

**Fig. 1.**
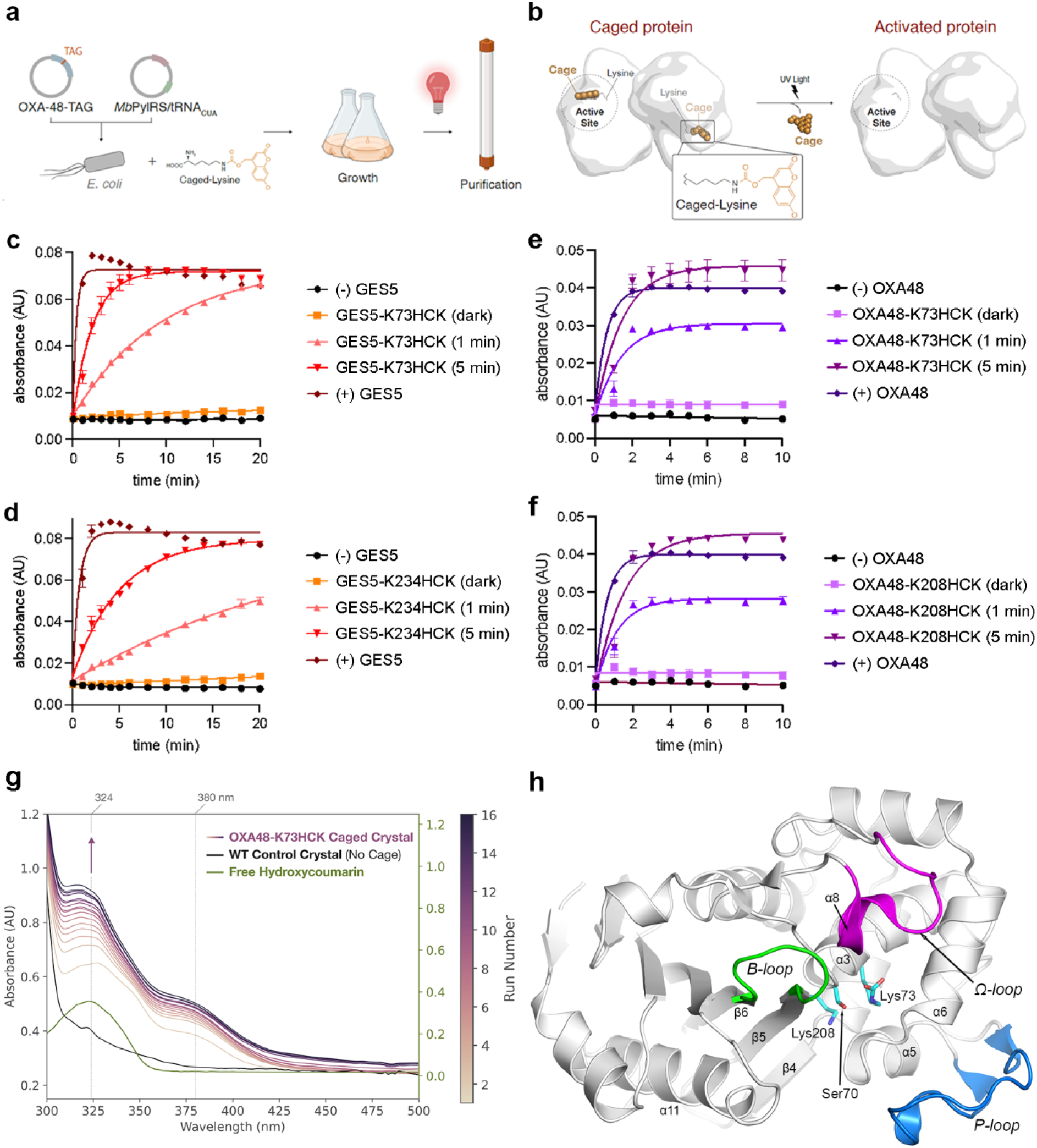
UAA mutagenesis to produce photocaged enzymes. (a) Schematic of UAA mutagenesis (see Supplementary Note 1) **(b)** *Release* workflow used to generate active enzymes. **(c-f)** Nitrocefin assays at 486 nm demonstrating caging and light-induced decaging of GES5-K73HCK (c), GES5-K234HCK (d), OXA48-K73HCK (e), and OXA48-K208HCK (f). Assays were monitored for 20 min (GES-5) or 10 min (OXA-48). Data were fit using a four-parameter variable slope model; error bars represent the standard deviation (n = 3) **(g)** UV decaging of an OXA48-K73HCK crystal shows a steady increase in a peak at 324 nm. The spectrum of free hydroxycoumarin shows a peak at approximately the same wavelength. **(h)** Ribbon representation of WT_A_ OXA-48 with selected secondary structure annotation. The P-loop is blue, the Ω-loop is pink, the B-loop is green, and residues Ser70, Lys73 and Lys208 are shown as cyan sticks.

We evaluated this strategy using serine β-lactamases, selecting the class A enzyme GES-5 and the class D enzyme OXA-48 as test systems. These enzymes are well suited due to their relatively simple architectures, well-defined active sites, extensively characterized catalytic mechanisms, and the availability of high-resolution crystal structures of substrate complexes^26-30^. These enzymes catalyze a two-step reaction comprising acylation of an active-site serine via a tetrahedral transition state intermediate, followed by deacylation to yield inactive product (Supplementary Fig. S2a) and support a straightforward spectrophotometric assay using nitrocefin. In addition to the catalytic serine, class A and class D β-lactamases contain two conserved lysine residues essential for catalysis (Supplementary Figs. S2b, S2c). These lysines were targeted for incorporation of hydroxycoumarin-caged lysine (HCK) (Supplementary Fig. S1c) via genetic code expansion: Lys73 and Lys234 in GES-5 (Ambler numbering), and Lys73 and Lys208 in OXA-48 (class D numbering^31,32^). Additional details are given in Supplementary Note 1.

Two GES-5 mutants (GES5-K73HCK and GES5-K234HCK) were generated, expressed and purified alongside wild-type (WT) enzyme. ESI–MS analysis confirmed incorporation of the photocaged amino acid, with observed mass shifts in close agreement with theoretical values (Supplementary Figs. S3a, S3b, S3c). Both mutants were inactive in nitrocefin assays prior to irradiation, but UV exposure (365 nm) restored activity, with partial recovery after short irradiation and full restoration after several minutes (Figs. 1c, 1d), consistent with efficient decaging. Similarly, two OXA-48 mutants (OXA48-K73HCK and OXA48-K208HCK) were produced and validated by ESI–MS (Supplementary Figs. S3d, S3e, S3f). These mutants were also inactive in the dark and regained activity upon UV irradiation, with shorter exposure yielding partial activity, indicative of incomplete decaging (Figs. 1e, 1f). Because class D β-lactamases require carboxylation of Lys73 for catalysis, decaging is expected to directly generate the catalytically competent residue, although reduced activity at shorter irradiation times may reflect incomplete decaging and/or loss of carboxylation. Decaging *in crystallo* was monitored by exposing an OXA48-K73HCK UV light, and recording spectra (Fig. 1g) using an *in situ* UV-visible microspectrophotometer at SSRL BL9-2^33,34^. The spectra showed a progressive increase in absorbance at 324 nm, which coincides with the λmax of free hydroxycoumarin (HC), indicating UV-induced cleavage of the photocage within the crystalline lattice. The time-dependent increase in signal is consistent with accumulation of the released HC photoproduct and provides direct spectroscopic evidence that decaging proceeds *in crystallo* under the illumination conditions used.

Crystallization trials were carried out for multiple constructs using a range of conditions previously established for the corresponding wild-type enzymes^27,28,35,36^. GES5 variants yielded only microcrystals unsuitable for diffraction experiments, and further optimization of the conditions for these constructs will be pursued in future experiments. OXA48-K73HCK (hereinafter HC73) produced large, well-formed crystals under conditions similar to WT OXA-48, and thus this variant was selected as the focus of subsequent structural characterization and decaging analyses. The WT and HC73 quaternary structures are shown in Supplementary Figs. S4a, S4b. Both enzymes crystallized with four monomers (A, B, C and D) in their asymmetric unit, arranged as a dimer of dimers (A/B and C/D). The structure of the WT monomer A (hereinafter WTA) is shown in Fig. 1h. The arrangement of each HC73 dimer (A/B or C/D) and their respective dimer interfaces are similar to the WT dimer. However, in contrast to the WT structure, the monomers which comprise the HC73 dimers exhibited conformational heterogeneity (Supplementary Note 2); monomers B and D have 15 residues missing from the Ω-loop while monomers A and C resemble their counterparts in WT. Since monomers C and D resemble monomers A and B in HC73, the remainder of this communication will consider only the A/B pair.

Structures determined under red-light conditions confirmed the presence of the intact hydroxycoumarin (HC) photocage. In one monomer (HC73A), the cage occupies the active site adjacent to the catalytic serine, with an occupancy of 70-80% (Fig. 2a). The HC photocage sits in the active site adjacent to the catalytic Ser70 and forms a steric barrier to incoming substrates. In the second monomer (HC73B), density corresponds exclusively to the caged lysine with full occupancy (Fig. 2b). The photocage, however, adopts an alternative conformation due to Ω-loop disorder, positioned outside the active site, overlapping the Ω-loop were it present. Molecular dynamics simulations (Supplementary Note 3, Supplementary Figs. S5, S6) support these observations, indicating that the cage remains stably confined within the active site of HC73A with continuous steric hindrance blocking of the catalytic serine, while exhibiting increased mobility and failure to enter the active site in HC73B.

**Fig. 2.**
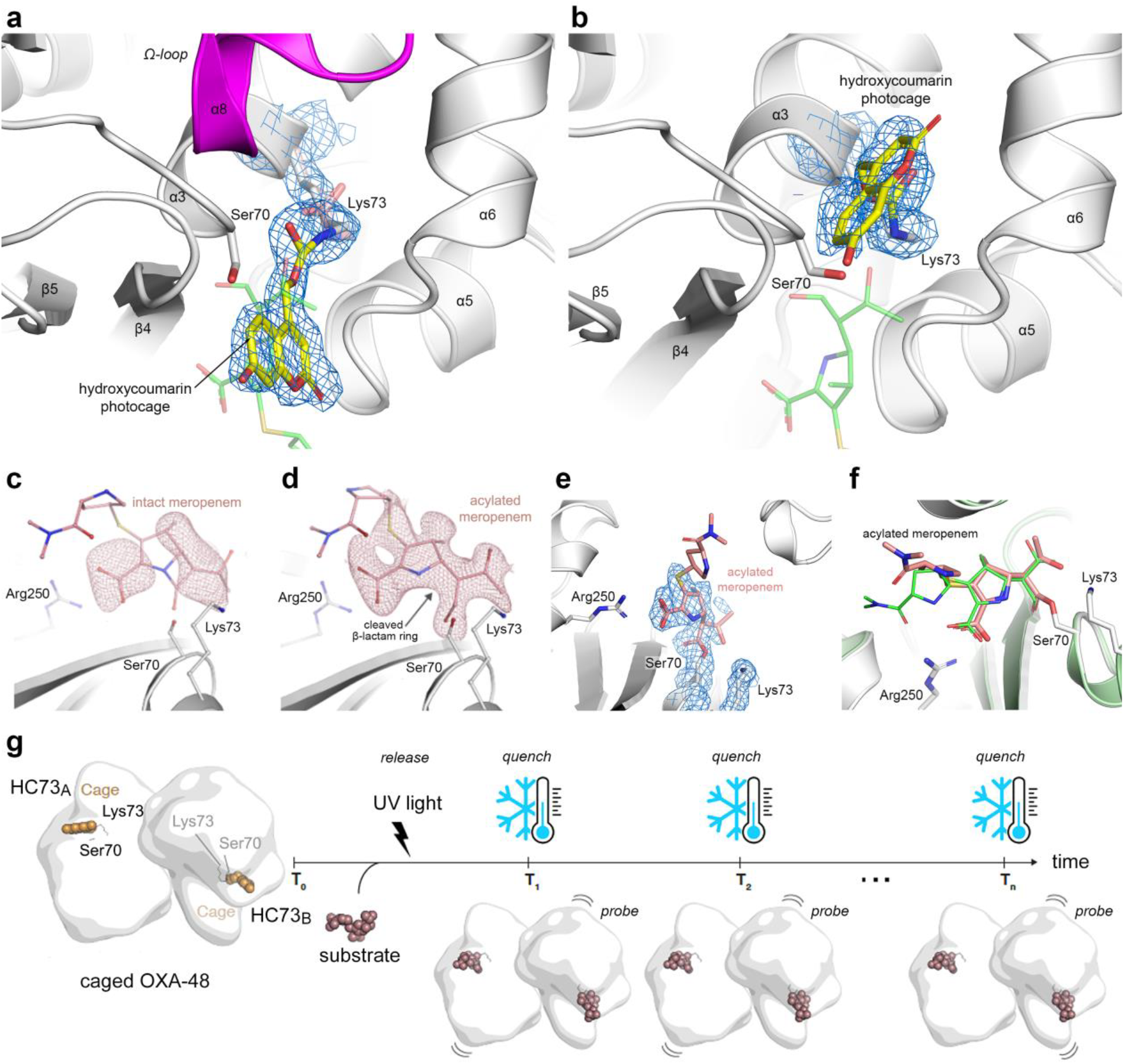
CAGE-TRX. Active sites of **(a)** HC73_A_ and **(b)** HC73_B_ showing 2*F*_*o*_*-F*_*c*_ density (blue mesh, 1σ) for the HCK. The position of acylated meropenem in WT OXA-48 is indicated by thin semi-transparent green sticks. **(c)** HC73A control structure with no UV exposure. Polder density (pink mesh, 3σ) is not consistent with either intact or acylated meropenem. **(d)** Polder density is consistent with β-lactam ring opening and acyl-enzyme formation. **(e)** Refinement of the resultant structure in panel **d** reveals strong 2*F*_*o*_*-F*_*c*_ density in the HC73_A_ active site for acylated meropenem and no detectable density for the HC photocage on Lys73. **(f)** Superposition of the partially refined HC73_A_-meropenem complex (white ribbons and pink meropenem) onto 7KHQ (pale green ribbons and green meropenem). **(g)** Schematic of the *release-quench-probe* CAGE-TRX workflow.

To assess functional consequences of HC photocaging *in crystallo*, meropenem was soaked into pre-formed photocaged crystals. The resultant HC73-meropenem structure also showed a clear dimorphism between the two monomers in each dimer. In HC73A, the active site was still occluded by the cage with no acylation of the catalytic serine observed, confirming that the enzyme remains inactive. Conversely, in HC73B, clear electron density consistent with an acyl-enzyme intermediate was observed (Supplementary Fig. S7), indicating that Ω-loop disorder permits substrate access and catalysis despite the presence of the caged residue (Supplementary Note 4). To confirm that UV illumination effectively released the HC photocage, experiment was undertaken where a HC73 crystal was irradiated with a UV flash and flash-cooled at 100 K. Structural analysis of the HC73A and HC73B active sites showed that in both cases the HC photocage was no longer attached to the Lys73 (Supplementary Figs. S8a, S8b).

For *release-quench-probe* CAGE-TRX experiments, photocaged crystals are soaked in a stabilizing solution containing substrates or inhibitors and mounted on the goniometer in a humid air stream. Crystal soaking can be performed immediately prior to mounting, or offline before the beamline experiments, since the enzymes have been rendered inactive by the photocaging. While in the humid stream, crystals are irradiated with a UV flash from a 1000 W Newport arc lamp (Supplementary Fig. S9a) to initiate decaging and displacement of the cage away from the active site (the *release* step). Reaction intermediates are subsequently trapped by rapid switching to a 100 K nitrogen stream using a mechanical cryo-trapping device built in-house^33^ (Supplementary Fig. S9b), which reproducibly transitions from the humid stream within 250 ms from triggering the device (the *quench* step). The timing of the cryo-switch relative to UV exposure defines the accessible time regime. Triggering the switch at or before UV exposure can capture short-lived intermediates on the millisecond timescale (∼25 -250 ms), whereas delayed switching enables observation of longer-lived states from hundreds of milliseconds to seconds or minutes. Following freeze-quenching, the cryo-cooled samples are dismounted using the SAM robot^37,38^ and stored in SSRL cassettes or Uni-pucks for subsequent data collection (the *probe* step). By eliminating the need for specialized caged substrates or rapid mixing approaches, CAGE-TRX provides a generalizable and experimentally accessible route for expanding TRX to a broader range of enzyme systems, with potential to support automated and remote synchrotron-based workflows.

As a proof-of-principle demonstration of CAGE-TRX for following catalysis catalysis *in crystallo*, photocaged HC73 crystals were soaked with meropenem under red light conditions and then subjected to a *release-quench-probe* workflow. Although preliminary, we were able to show an increase in meropenem acyl-enzyme occupancy in HC73A following reaction initiation. A control structure collected prior to UV exposure (*quench-probe* only; Fig. 2c) exhibited weak electron density near catalytic Ser70; however, this density was not consistent with either intact meropenem (the Michaelis complex) or an acylated species and may reflect partial hydrolysis of the HC photocage. In contrast, a structure from a crystal subjected to the complete *release-quench-probe* workflow showed substantial accumulation of electron density in HC73A consistent with formation of an acylated meropenem intermediate (Fig. 2d). An acyl-enzyme complex was modeled and partially refined, revealing strong electron density supporting increased occupancy of acylated meropenem (Fig. 2e). Comparison of the HC73A acyl-enzyme complexes with a known OXA-48-meropenem acyl-enzyme complex (PDB code 7KHQ) shows that the acylated carbapenems are bound in almost identical fashion (Fig. 2f). Together, these results indicate that photodecaging restores catalytic activity *in crystallo* and enables capture of a *bona fide* reaction intermediate.

Together, these findings establish a proof-of-principle for controlling enzyme activity in protein crystals using photocaged amino acids. We develop and validate CAGE-TRX, a robust workflow for generating catalytically silent enzymes that can be activated on demand *in crystallo*, enabling synchronized initiation of catalysis throughout the crystal (Fig. 2g). Structural analysis confirms that caging blocks substrate access and prevents catalysis, while photolysis restores active-site accessibility, restores activity and allows progression through the reaction coordinate. In the OXA-48 system, this approach enables time-resolved trapping of an authentic acyl-enzyme intermediate following reaction with a clinically relevant β-lactam substrate, and may even allow for the capture of the Michaelis complex and, more importantly, the elusive tetrahedral transition state intermediate. Given the high degree of sequence, structural, and mechanistic conservation across serine β-lactamases, this strategy is likely broadly applicable for probing antibiotic turnover, inhibition, and resistance amongst these clinically important enzymes.

Photocaging of amino acids provides a modular and generalizable approach for time-resolved structural biology. Established chemistries enable caging of multiple residue types including lysine, tyrosine, cysteine, serine histidine, phenylalanine, glutamate and aspartate^25^, many of which are frequently found in enzyme active sites^39^. Combined with mature genetic code expansion technologies^23,25,40,41^, this approach offers broad access to enzyme systems that are perhaps not readily amenable to conventional TRX approaches. While photocaged substrates exist for some systems, their development is often enzyme-specific and synthetically demanding^18^. CAGE-TRX decouples reaction triggering from substrate design, provides a substrate-independent route to light-controlled catalysis *in crystallo*, and expands the scope of time-resolved crystallography to a wider range of enzymatic systems. By enabling uniform, light-triggered activation in the presence of substrate within solvent channels, CAGE-TRX minimizes spatial and temporal heterogeneity associated with diffusion-based methods^42,43^ and is compatible with both cryogenic trapping and RT serial synchrotron crystallography. As such, it expands the experimental toolkit for probing enzyme mechanism, inhibition and antibiotic resistance, and provides a broadly applicable framework for capturing transient states in enzymatic reactions.

## Methods

### UAA mutagenesis of GES-5 and OXA-48

The expression constructs pBAD-GES5-pylT and pBAD-OXA48-pylT were generated via Gibson assembly^44^. The GES5 and OXA48 fragments were PCR amplified from pET24a-GES5 and pET24a-OXA48 using primers 1/2 and 3/4, respectively, at which point an N-terminal His-tag was also introduced for protein purification. The pBAD-pylT backbone was generated via restriction enzyme digest of pBAD-sfGFP-pylT (2000 ng) using NcoI (NEB R013S) and NdeI (NEB R0111S) and subsequent gel purification. The expression constructs pBAD-GES5-K73TAG-pylT, pBAD-GES5-K234TAG-pylT, pBAD-OXA48-K73TAG-pylT, pBAD-OXA48-K208-TAG-pylT were generated from the corresponding pBAD-beta-lactamase construct via site-directed mutagenesis using a QuikChange design. Primers were designed using the Agilent QuikChange Primer Design Tool online and PCR was performed according to the Agilent QuikChange manual. Primers 5/6 were used for pBAD-GES5-K73TAG-pylT, primers 7/8 were used for pBAD-GES5-K234TAG-pylT, primers 9/10 were used for pBAD-OXA48-K73TAG-pylT, and primers 11/12 were used for pBAD-OXA48-K208TAG-pylT (Supplementary Table S1).

### Expression of wildtype enzymes

The pBAD-GES5-pylT or pBAD-OXA48-pylT plasmids (50 ng) were transformed into chemically-competent Top10 cells and plated on LB-agar (10 mL, + 25 µg/mL tetracycline). A single colony for each was grown in LB broth (5 mL, + 25 µg/mL tetracycline) overnight at 37 °C with 250 rpm shaking to generate a saturated starter culture. In a 500 mL Erlenmeyer flask, LB broth (100 mL, + 25 µg/mL tetracycline) was inoculated with 2 mL of the saturated starter culture. The expression cultures were grown at 37 °C with 250 rpm shaking until the OD600 reached 0.6, then protein expression was induced using 400 µL of 20% arabinose for a final concentration of 0.08% arabinose. Expression was performed at 37 °C with 250 rpm shaking for ∼5 hours, until an OD600 of 1.8 was reached. At this time, cells were harvested by centrifugation at 4500 rpm for 10 minutes at 4 °C in 50 mL conical tubes and the supernatant was discarded. Cell pellets were stored at –80 °C until the next day. Cell pellets for each protein were gently resuspended and combined in a total of 10 mL of bacterial lysis buffer [300 mM NaCl, 45 mM Na2HPO4, 5 mM NaH2PO4, 3 mM NaOH, 10 mM imidazole, 1 mg/mL lysozyme, 1 µL/mL protease inhibitor cocktail (Sigma P8849), 0.01% (v/v) Triton X-100], then incubated on ice for 20 minutes. Cell resuspensions were sonicated at 60% amplitude for 3 minutes with cycles of 10 seconds on, and 10 seconds off. Cell lysates were centrifuged at 4 °C at 4500 rpm for 10 minutes to pellet cellular debris. The soluble protein fractions were transferred to 15 mL conical tubes for Ni-NTA resin purification.

First, the Ni-NTA resin (G-Biosciences 786939) was washed three times with lysis buffer to remove the 20% ethanol solution. After washing, the resin (125 µL) was resuspended in an equal volume of lysis buffer (125 µL), then added to the soluble protein fraction. This solution was incubated at 4 °C for 2 hours on a multi-directional rocker. After this time, the resin was collected by centrifugation at 4 °C for 10 minutes at 800 g, and the supernatant was removed. The resin was washed 3 times using 300 µL of bacterial wash buffer [9 mL bacterial lysis buffer + 1 mL bacterial elution buffer], removing the supernatant between each step by centrifugation at 4 °C for 2 minutes at 800 g. Protein was eluted from the resin 4 times using 200 µL of bacterial elution buffer [50 mM NaH2PO4, 300 mM NaCl, 250 mM imidazole, 20 mM NaOH] and the purified proteins were visualized via SDS PAGE. Protein concentrations were determined via Bradford assay.

### Expression and purification of photocaged enzymes

Expression and purification were performed as described above but starting with a double transformation of the appropriate pBAD-GES5-TAG-pylT and pBK-HCKRS. Approximately 15 minutes prior to induction, HCK (100 mM in DMSO) was added to a final concentration of 1 mM. Expression was performed at 37 °C with 250 rpm shaking for ∼5 hours, until an OD600 of 1.8 was reached. At this time, cells were harvested by centrifugation at 4500 rpm for 10 minutes at 4 °C in 50 mL conical tubes and the supernatant was discarded.

Since preliminary crystallization trials suggested that the presence of the N-terminal His6-tag was interfering with crystal formation, the tag was removed prior to shipping the enzymes to SSRL. Cell pellets were stored at –80 °C. The pellets were resuspended in 25mM Hepes, pH7.5 buffer and disrupted by sonication at 4 °C. The suspension was supplemented with NaCl to 200mM and centrifuged at 100,000g for 1hr. The supernatant with the soluble proteins was adjusted to 20mM imidazole and passed through the HiTrap Chelating HP column (Cytiva) which was charged with Ni-sulfate and equilibrated with binding buffer (25mM Hepes, pH7.5, 300mM NaCl, 20mM imidazole) according to instructions. The column was washed with 5 column volume of binding buffer and the N-terminus His6-tagged protein was eluted with discontinuous gradient of the binding buffer containing 20, 50, 100, and 250mM imidazole (3 column volumes each). Fractions were analyzed on SDS-PAGE, and those containing the target protein were combined and concentrated using the 10K cut off concentrator. Imidazole was removed by dialysis in a buffer containing 25mM Hepes, pH 7.5 and 150mM NaCl. The N-terminus His6-tag was cleaved by the His6-tagged tobacco etch virus (TEV) protease during overnight incubation at 15°C with gentle rotation. The solution was centrifuged at 20000g for 10 minutes and the supernatant was adjusted to 300 mM NaCl and passed through the Ni^2+^-column equilibrated with the binding buffer. Photocaged proteins were collected in the flow through fractions obtained by washing the column with the binding buffer. Fractions containing the target protein were combined, concentrated and dialyzed against 25mM Hepes, pH 7.5. For protein crystallization the purified enzymes were further concentrated and stored at 4 °C. All steps of the protocol were performed under dark or red-light conditions to avoid premature decaging.

### Crystallization, data collection and structure solution

The wild-type (WT) OXA-48 enzyme was crystallized from published conditions (100 mM HEPES pH 7.5, 8% 1-butanol, 10% PEG8000)^26^ at 10°C. The crystals were transferred to a cryo-buffer comprising the crystallization reservoir solution augmented with 30% glycerol. Diffraction data were collected to approximately 1.6 Å resolution at SSRL beamline BL12-2 with X-rays of wavelength 0.97965 Å (12658 eV) on a Pilatus 6M PAD detector. The crystals indexed in space group P21, with unit cell parameters a, b, c = 58.6 Å, 106.5 Å, 94.7 Å and β = 107.4°. The data were processed and scaled with XDS^45^ and AIMLESS^46^.

HC73 was crystallized under multiple conditions known to give crystals^28^, with crystals appearing within 1-2 days under the same conditions as WT. All manipulations of the mutant enzyme were carried out inside a cold room at 4°C with the main lights off, using red light from a 4.5 W Sylvania LED bulb. The microscope stage was covered with a red filter to eliminate short wavelength light including UV. After crystallization drop setup, the trays were wrapped in aluminum foil and stored under zero-light conditions at 10°C. Crystals were flash-cooled in the same cryo-buffer as the WT crystals, again in the cold room under red light to minimize UV exposure, and stored in an SSRL cassette^37^ in the dark. The crystals were rods at least 250 μm in length, 40-50 μm across and 20 μm thick, and resembled those observed for the WT enzyme. Despite the identical crystallization conditions and crystal morphology, the HC73 crystals adopted a different space group, P212121, with unit cell parameters of *a, b, c* = 90.8 Å,

106.4 Å, 125.5 Å. A data set extending to 1.75 Å resolution was collected on BL9-2 with X-rays of wavelength 0.97965 Å on a Pilatus-6M PAD detector, and processed and scaled with XDS^45^ and AIMLESS^46^. Data collection statistics are given in Supplementary Table S2.

To test the *in-crystallo* activity of HC73, several pre-formed crystals were transferred into cryo-buffer augmented with 10 mM meropenem and incubated at 4°C in the dark for 2-3 minutes. The crystals were then flash cooled in liquid nitrogen and stored in the dark in an SSRL cassette for subsequent data collection. Data were collected at SSRL beamline BL9-2 with X-rays of wavelength 0.97965 Å on a Pilatus-6M PAD detector, and the images were processed and scaled with XDS^45^ and AIMLESS^46^.

For preliminary TRX experiments, following de-caging and freeze-trapping (described below), the HC73 crystals stored inside an SSRL cassette were robotically mounted for subsequent diffraction data collection at SSRL BL9-2 and BL12-2. All data sets were processed and scaled with XDS^45^ and AIMLESS^46^.

### *In crystallo TR* decaging and freeze-quenching

Sample freeze-trapping was carried out using a mechanized freeze-trapping system (Supplementary Fig. S8b) developed in-house by SSRL-SMB, and incorporated into the SSRL BL7-1 experimental hutch, a beamline primarily used for in-house developments. For rapid freeze-trapping of intermediate states within crystals, the system uses a rotary actuator to rapidly swap between a humid sample environment (produced by an Arinax HC-Lab Humidity Controller) and a cryogenic sample environment produced by a cold nitrogen gas stream (100 K, Oxford Cryosystems 800 cryojet), a transition which takes approximately 250 ms. Two programmable Berkeley Nucleonics Corporation Model 725 programmable logic and timing controllers are incorporated into the freeze-trapping system. These are used for coordinating and synchronizing system operation, including motion of the rotary actuator, gas line divertors, and a Thorlabs SHB025T high-speed optical shutter, with the readout of light and positional sensors for timing feedback and verification. Light produced by a 1000 W Newport arc flash lamp passes through an IR filter and the fast shutter before entering a 5X UV-objective, to produce a ∼1.4 mm (FWHM) light spot at the sample/crystal position (Supplementary Fig. S8a). Using this system, crystals could be exposed to pulses of UV light and then flash cooled. By initiating the flash cooling process (transition between the humid and cryo-nozzle) before initiating a light pulse, fast reaction intermediates (>5 ms) can be freeze-trapped. Prior to freeze-quenching, HC73 crystals were soaked in cryo-buffer augmented with 10 mM meropenem and hand-mounted onto a Huber goniometer in standard nylon loops (Hampton Research). The humidity control device positioned co-linear with the phi axis of the goniometer was used to maintain the crystals at room temperature. A WT crystal was used to verify the tabulated relative humidity value (92%) for a solution containing 100 mM HEPES, 8% 1-butanol, 10% PEG8000 and 30% glycerol prior to mounting the HC73 crystals on the goniometer and humid stream environment.

To minimize UV exposure from background sources, all lights within the BL7-1 hutch were either turned off or covered with a red filter. Since the hutch door is not permitted to be closed with personnel inside due to radiation safety protocols, a blackout curtain was installed on a rail just inside the door. A 4.5 W red LED was used for additional lighting, and a microscope with a red filter was used for crystal manipulation.

A series of experiments were performed with varying UV flash durations and delays between initiating the UV flash and freeze-trapping, to create timepoints along the TRX trajectory. For each experiment, we recorded the UV flash duration and the delay before the crystal was positioned in the cooling stream. After freeze-trapping, the crystals remained in the 100 K cryostream environment until they were dismounted using the SAM robot^37,38^ and stored in SSRL cassettes for subsequent data collection.

## Acknowledgements

This work was supported, in part, by DOE-BER BRaVE, a DOE LDRD award to C.A.S., and the National Institutes of Health through grant R01AI175067 to A.D., and grant R01AI155723 to S.B.V. Parts of this research were carried out at the Stanford Synchrotron Radiation Lightsource (SSRL), a national user facility operated by Stanford University on behalf of the U.S. Department of Energy. Use of SSRL is supported by the U.S. Department of Energy, Office of Science, Office of Basic Energy Sciences under Contract No. DE-AC02-76SF00515. The SSRL Structural Molecular Biology Program is supported by the DOE Office of Biological and Environmental Research, and by the National Institutes of Health, National Institute of General Medical Sciences (P30GM133894). The contents of this publication are solely the responsibility of the authors and do not necessarily represent the official views of NIGMS or NIH.

## Author contributions

C.A.S., A.D., G.C. and A.E.C. developed the underlying scientific ideas for CAGE-TRX. C.J., M.C. and N.P. performed the UUA mutagenesis, enzyme expression and purification, along with activity assays. M.T. and S.B.V. carried out purification of OXA mutants for crystallization. C.A.S. and A.O.M. grew crystals of wild-type and mutant GES and OXA enzymes. S.C. and A.E.C. designed, manufactured, installed and tested all *release-quench-probe* workflow components at SSRL BL7-1. A.O.M., C.A.S., S.C. and S.R. performed all *release-quench-probe* work. C.A.S. and A.E.C. wrote the draft of the manuscript. All authors contributed to editing the manuscript. Figures were created by A.O.M. and C.A.S.

## Notes

### Competing Interest Statement

The authors have declared no competing interest.

