## Supplementary Tables, Figures and Notes for "CAGE-TRX expands the scope of time-resolved crystallography through genetically encoded active-site photocaging"

**Table S1.** List of primers used for plasmid construction (shown 5' to 3'). Base mutations are indicated by capitalization.

| PRIMER | SEQUENCE |
|---|---|
| **P1** | aacaggaggaattaaccatggggcatcatcatcatcatcatgaaaatctttattttcaaggtgcgtctgagaaa |
| **P2** | ggcccaagcttcgaattcccatatgttatttatcggtggacaggattaactgcgt |
| **P3** | aacaggaggaattaaccatggggcatcatcatcatcatcatgaaaatctttattttcaaggtgcgaaagaatgg |
| **P4** | ggcccaagcttcgaattcccatatgttacggaatgatttttttcttgtttcagga |
| **P5** | atgtgtagtacctttTAGtttccgttagcggcc |
| **P6** | ggccgctaacggaaaCTAaaaggtactacacat |
| **P7** | gctggctggtttattgctgatTAGtctggagccggtgag |
| **P8** | ctcaccggctccagaCTAatcagcaataaaccagccagc |
| **P9** | ccggcaagcacgttcTAGattccgaattctctg |
| **P10** | cagagaattcggaatCTAgaacgtgcttgccgg |
| **P11** | tatattatccgtgcgTAGaccggttacagcacg |
| **P12** | cgtgctgtaaccggtCTAcgcacggataatata |

**Table S2. OXA-48 HC73 data collection and refinement statistics**

| | | HC73 |
|---|---|---|
| *Data Collection* | | |
| Unit cell, a, b, c (Å) | | 90.79, 106.36, 125.48 |
| Resolution (Å) | | 39.6-1.75 (1.78-1.75)[a] |
| Reflections | - observed | 1633569 |
| | - unique | 122683 (5957) |
| $R_{meas}$[b] | | 0.127 (1.93) |
| $R_{pim}$[c] | | 0.034 (0.585) |
| $I$ / $\sigma_I$ | | 12.2 (1.2) |
| Completeness (%) | | 99.9 (99.2) |
| CC½[d] | | 0.999 (0.78) |
| Average multiplicity | | 13.3 (10.4) |
| Wilson B ($Å^2$) | | 24.3 |
| *Refinement* | | |
| $R_{work}$ / $R_{free}$[e] | | 0.2254 / 0.2521 |
| Reflections, work/free | | 116372 / 6165 |
| Number of atoms | - protein | 7745 |
| | - water | 563 |
| | - ligands | 64 |
| B-factors ($Å^2$) | - protein | 26.5 |
| | - water | 32.5 |
| | - ligands | 40.3 |
| *rmsd*s | - bond lengths (Å) | 0.002 |
| | - bond angles (°) | 0.91 |
| Ramachandran plot[f] | - favored (%) | 98.4 |
| | - outliers | 0 |
| Molprobity Score[g] | | 0.83 (100$^{th}$ percentile) |
| Molprobity Clashscore[g] | | 1.17 (99$^{th}$ percentile) |

[a] Numbers in parentheses refer to the highest resolution shell.
[b] $R_{meas}$ is the redundancy-independent merging R factor.[1]
[c] $R_{pim}$ is the precision-indicating merging R factor.[1]
[d] Correlation between intensities from random half-sets of data.[2]
[e] $R_{free}$ was calculated using a test set comprising 5% of the data.
[f] Calculated with the program MOLPROBITY.[3]

### Supplementary Figures

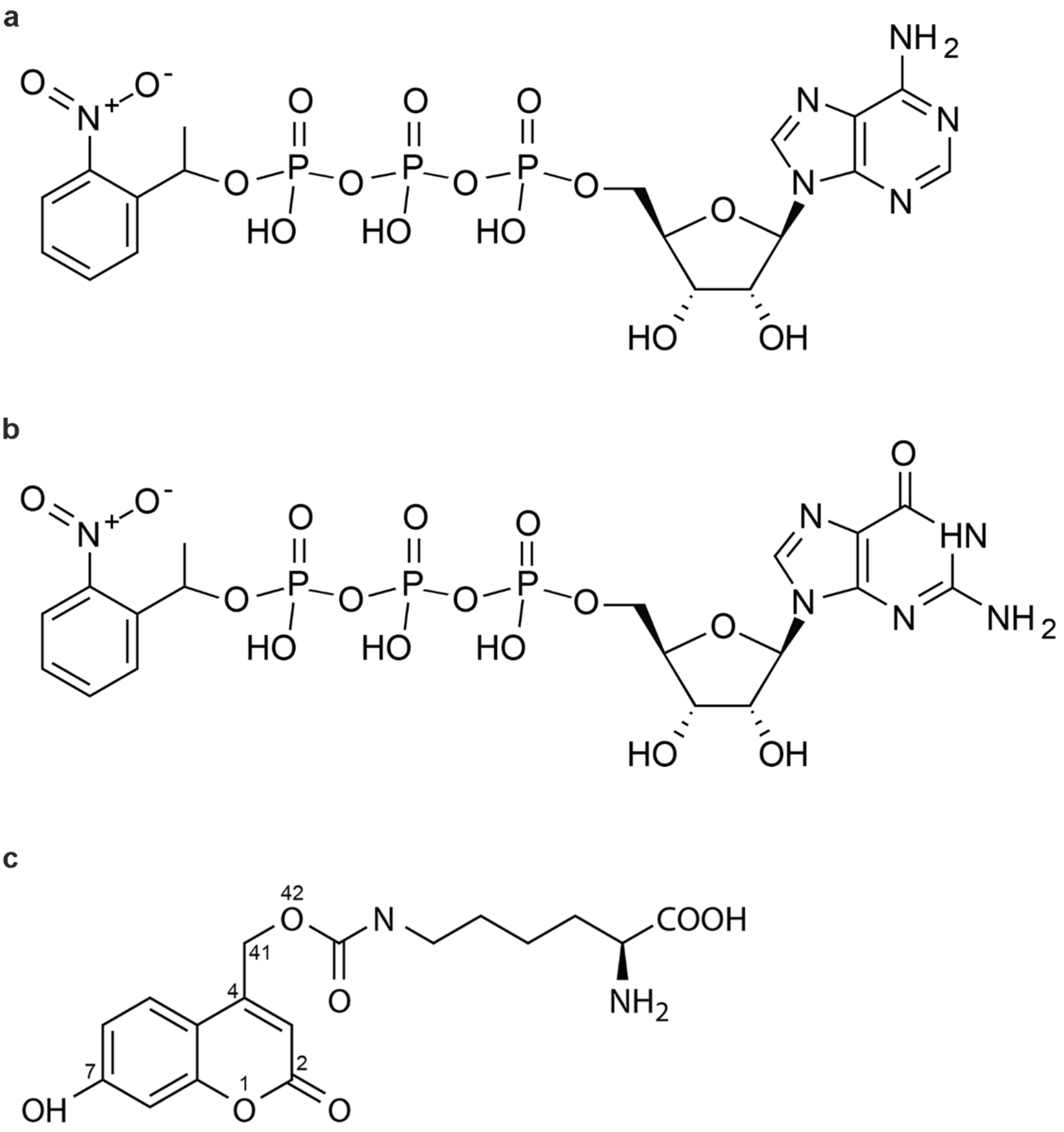


**Supplementary Fig. 1. Photocaged compounds. (a)** NPE-caged ATP. **(b)** NPE-caged GTP. **(c)** hydroxycoumarin-lysine (HCK).

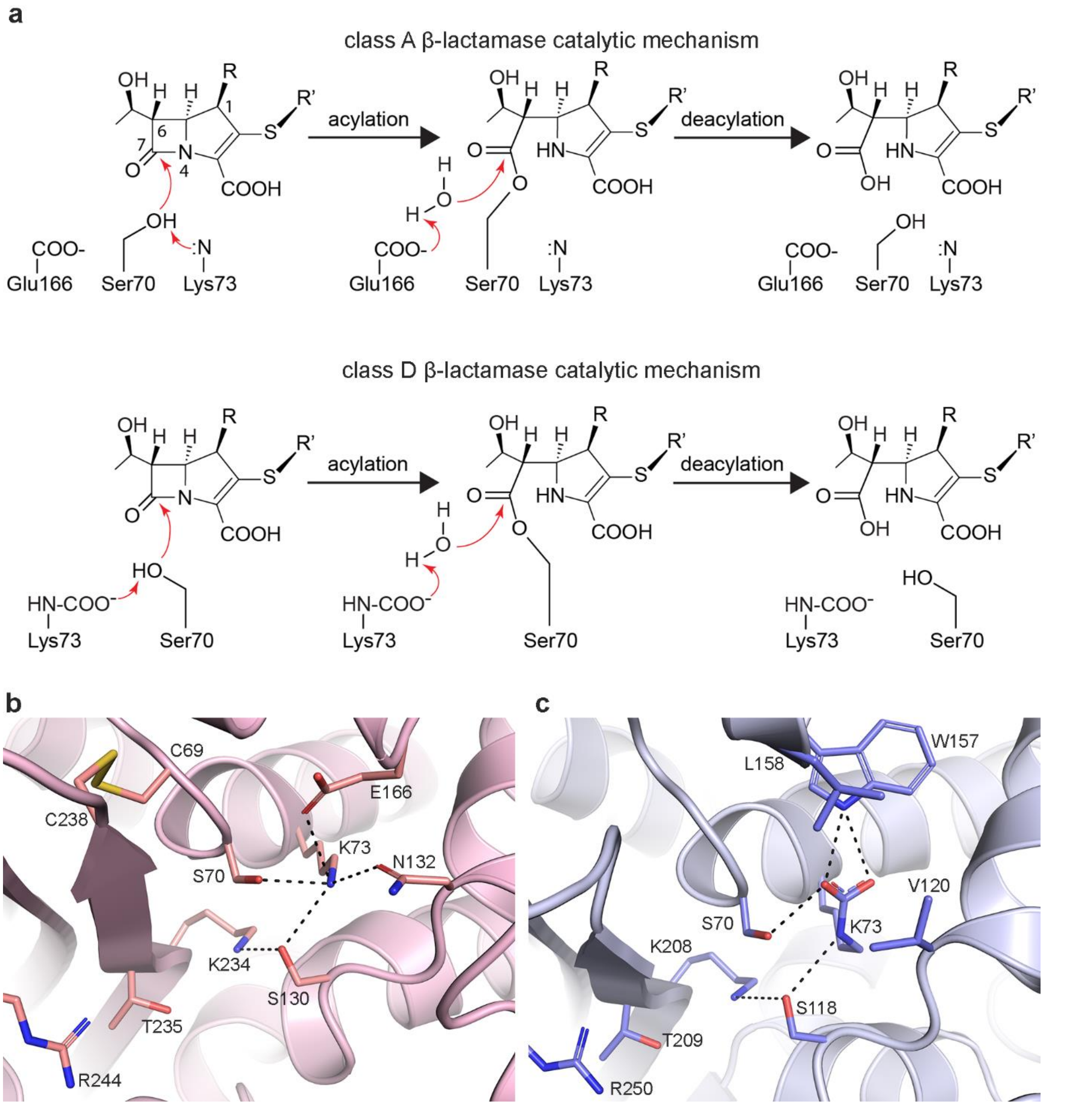


**Supplementary Fig. 2. Class A and class D β-lactamases. (a)** Generalized catalytic mechanisms of the class A and class D β-lactamases. Hydrolysis of a generic carbapenem is shown. The R group at position 1 is either hydrogen in imipenem or a methyl group in other carbapenems, including meropenem. The R' tail group is variable depending on the carbapenem. **(b)** Active site of the class A carbapenemase GES-5 (4GNU). **(c)** Active site of the class D β-lactamase OXA-14 (7L5R). The hydrogen bonding connections between Ser70, Lys73 and Ser130 (class A) or Ser118 (class D) create a proton shuttle necessary for acylation.

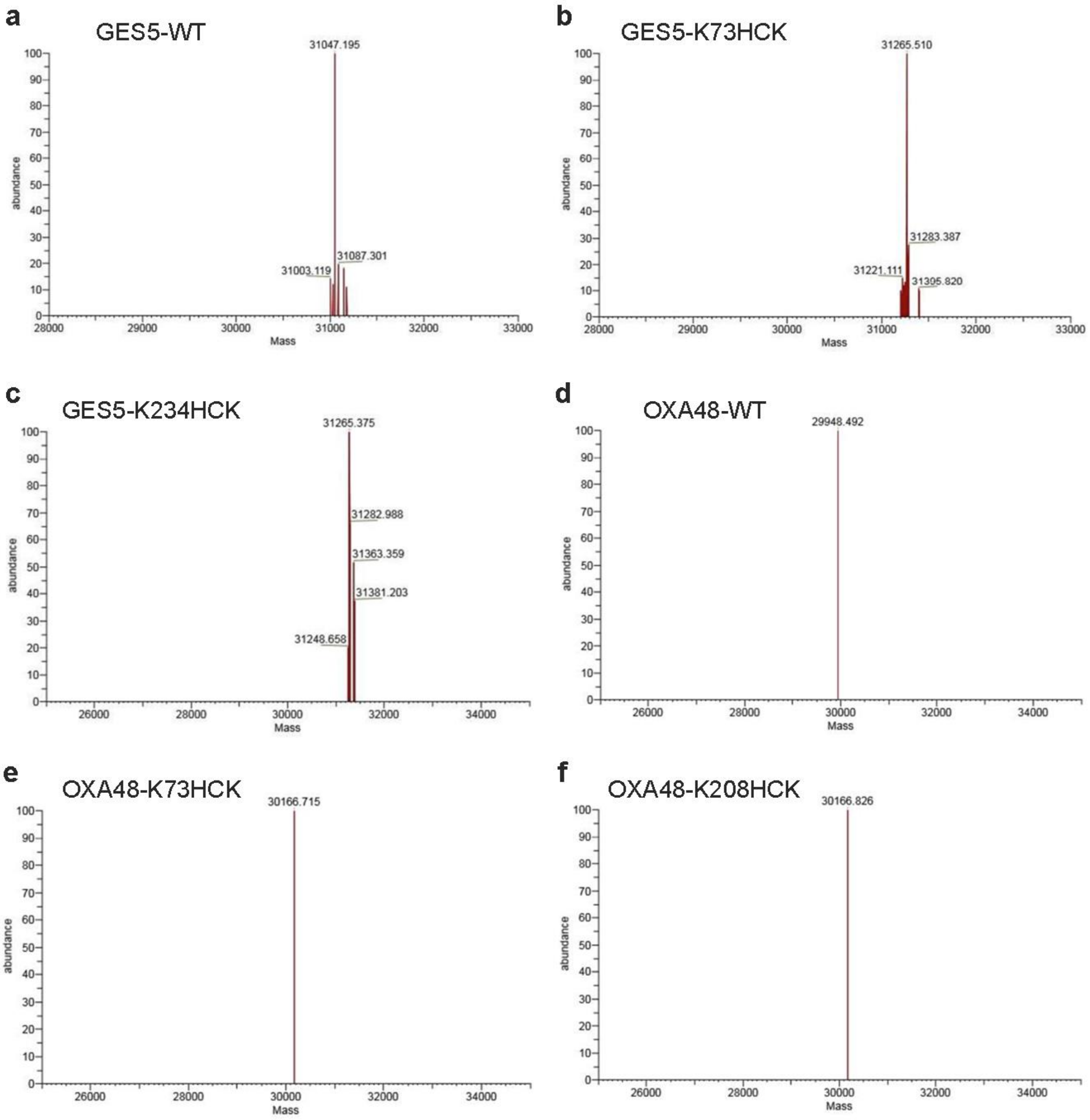


**Supplementary Fig. 3. Mass spectra of GES-5 and OXA-48 enzymes and mutants. (a)** Wildtype GES5 (31047.195 Da). **(b)** K73HCK (31265.510 Da, Δ 218.315 Da). **(c)** K234HCK (31265.375 Da, Δ 218.180 Da). Both mutants showed the expected mass shift (Δ 218.021 Da) corresponding to the replacement of lysine with HCK in GES-5. **(d)** Wildtype OXA48 (29948.492 Da). **(e)** K73HCK (30166.715 Da, Δ 218.223 Da). **(f)** K208HCK (30166.826 Da, Δ 218.334 Da). Both mutants showed the expected mass shift (Δ 218.021 Da) corresponding to the incorporation of HCK into OXA-48.

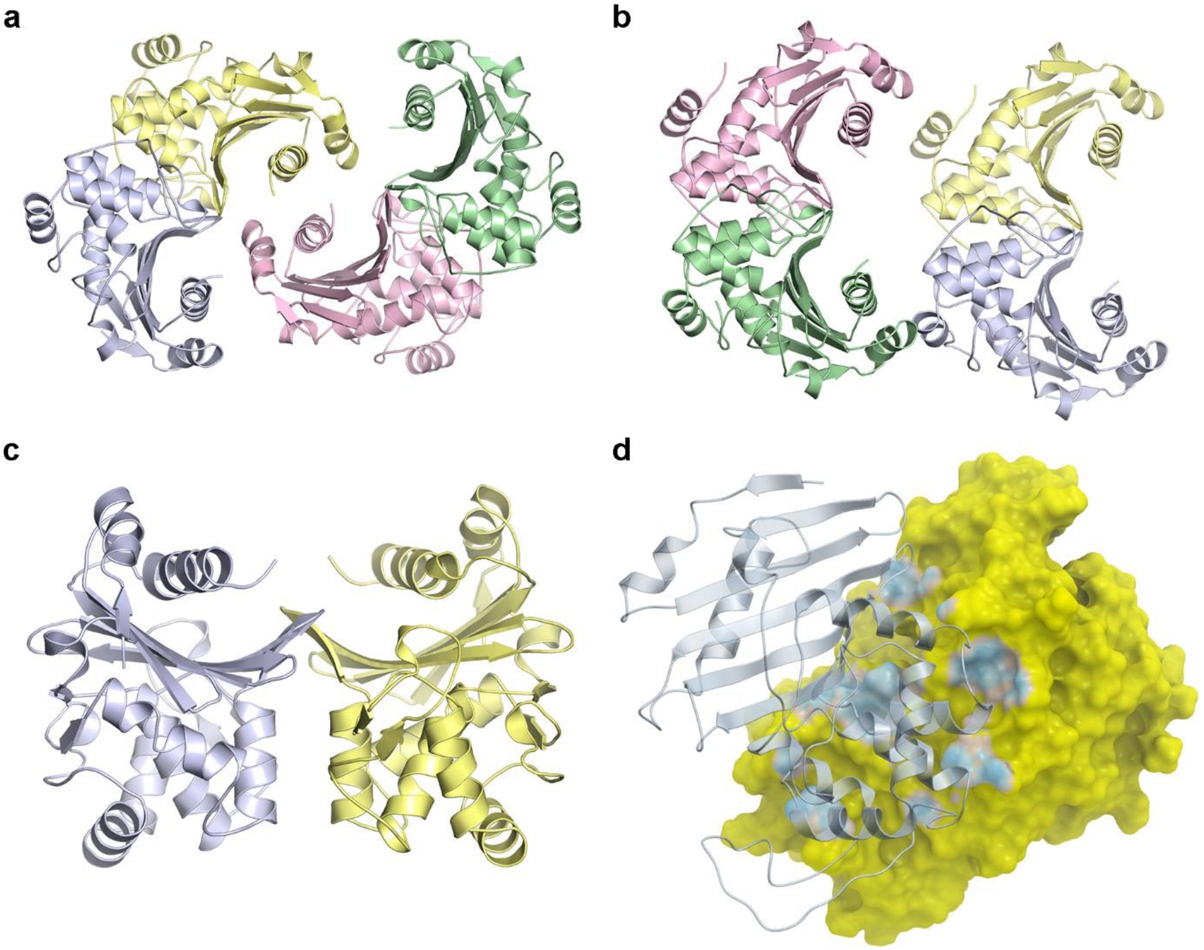


**Supplementary Fig. 4. Quaternary structure WT and HC73. (a)** The asymmetric unit of WT. **(b)** The asymmetric unit of HC73. **(c)** The WT A/B dimer, $WT_A$ (yellow) and $WT_B$ (light blue). **(d)** The WT A/B dimer interface. $WT_A$ is shown as a yellow molecular surface and $WT_B$ as light blue ribbons.

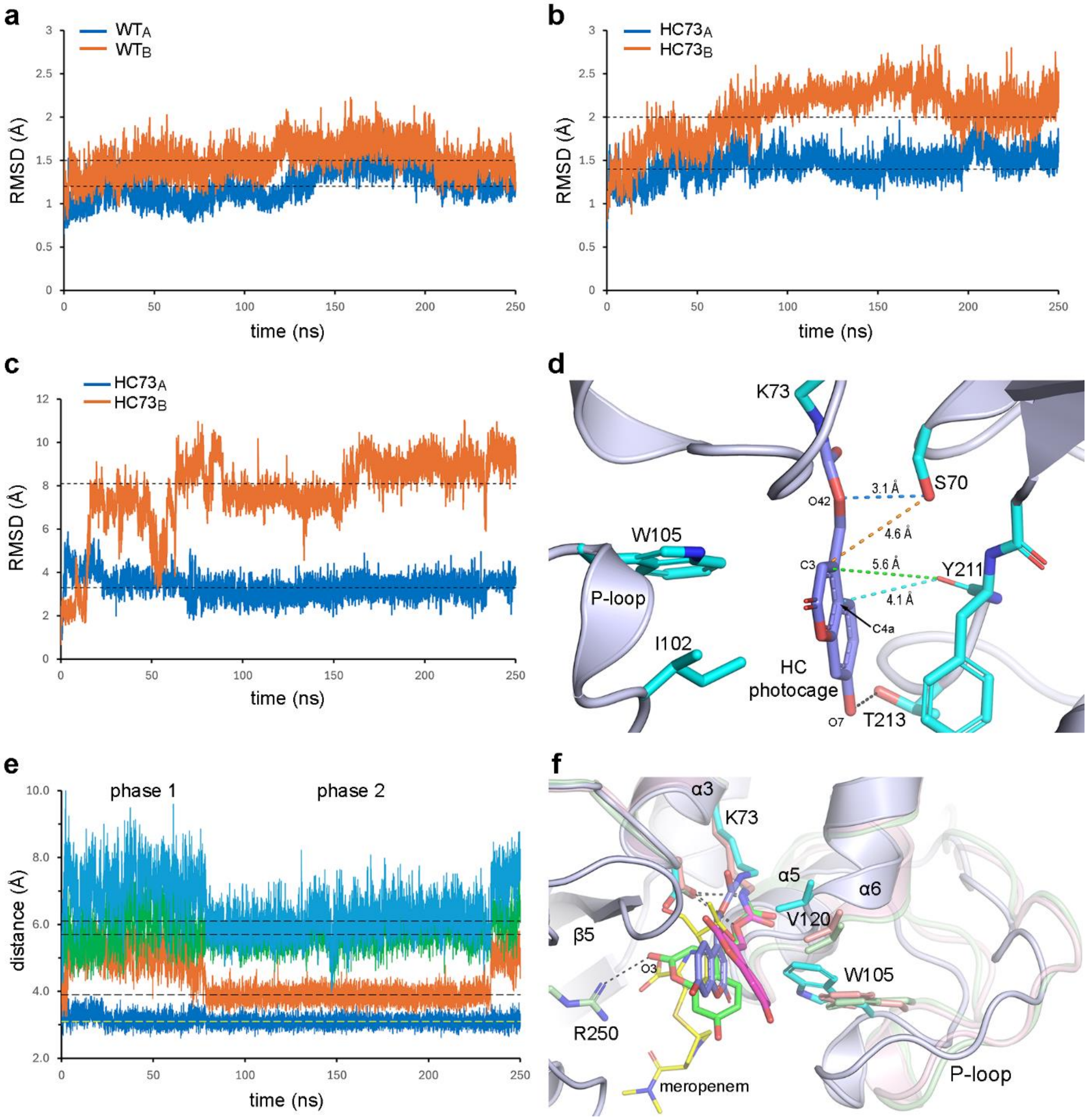


**Supplementary Fig. 5. MD simulation of HC73. (a)** Plot of RMS deviation (RMSD) for $WT_A$ (blue) and $WT_B$ (orange) protein atoms. The average RMSDs are indicated by black dashed lines; 1.2 Å for $WT_A$ and 1.5 Å for $WT_B$. **(b)** Plot of RMSD for $HC73_A$ (blue) and $HC73_B$ (orange) protein atoms. The average RMSDs are 1.4 Å for $HC73_A$ and 2.0 Å for $HC73_B$. **(c)** Plot of the RMSD of the HC photocage in $HC73_A$ (blue) and $HC73_B$ (orange) during a 250 ns molecular dynamics simulation using Desmond. The RMSDs were calculated for each 25 ps time point using the structure at $t = 0$ as the reference. **(d)** The $HC73_A$ active site showing four distances between the HC photocage and the protein used to monitor the position of the photocage. The distances shown are for the reference time point prior to any simulated motion of the HC photocage. **(e)** Plot of the four distances over the MD trajectory. The average distances are shown as dashed lines. Ser70Oγ – O42 (blue, 3.1 Å average), Ser70Oγ – C3 (orange, 3.9 Å), Tyr211O – C3 (green, 5.7 Å) and Tyr211O – C4a (cyan, 6.1 Å). **(f)** Superposition of three timepoints from the MD trajectory, $t = 0$ (light blue ribbons, cyan sticks, blue HC photocage), $t = 50$ ns (pink semi-transparent ribbons, pink sticks and magenta HC photocage) and $t = 200$ ns (green semi-transparent ribbons, green sticks and HC photocage). The position of acylated meropenem from 6PT1 is shown as thin yellow sticks.

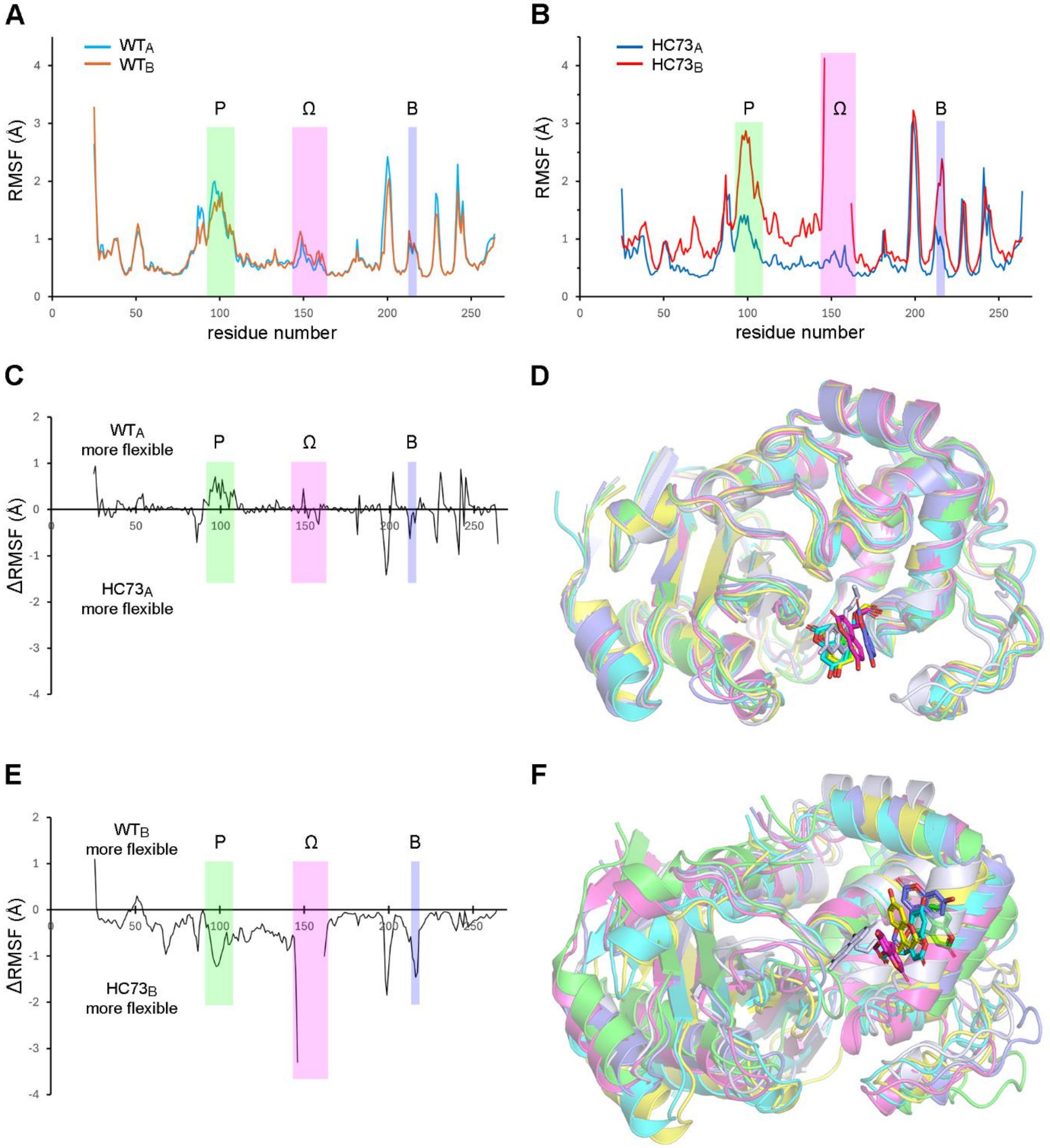


**Supplementary Fig. S6. Structural fluctuations in HC73. (A)** Plot of RMS fluctuations (RMSF) as a function of residue number for both WT monomers. **(B)** Plot of RMSF for both HC73 monomers. These plots allow identification of regions within the enzymes that exhibit more flexibility (high RMSF) or rigidity (low RMSF). **(C)** RMSF difference plot calculated as WTA – HC73A to highlight regions of increased flexibility in the two monomers. **(D)** Superposition of frames at five timepoints from the HC73A MD trajectory (*t* = 50 ns, magenta; *t* = 100 ns, cyan; *t* = 150 ns, yellow; *t* = 200 ns, green; *t* = 250 ns, blue) onto the HC73A crystal structure (light blue). The position of the HC photocage at each timepoint are indicated in the same colors. **(E)** RMSF difference plot calculated as WTB – HC73B. **(F)** Superposition of frames at five timepoints from the $HC73_B$ MD trajectory (*t* = 50 ns, magenta; *t* = 100 ns, cyan; *t* = 150 ns, yellow; *t* = 200 ns, green; *t* = 250 ns, blue) onto the $HC73_B$ crystal structure (light blue).

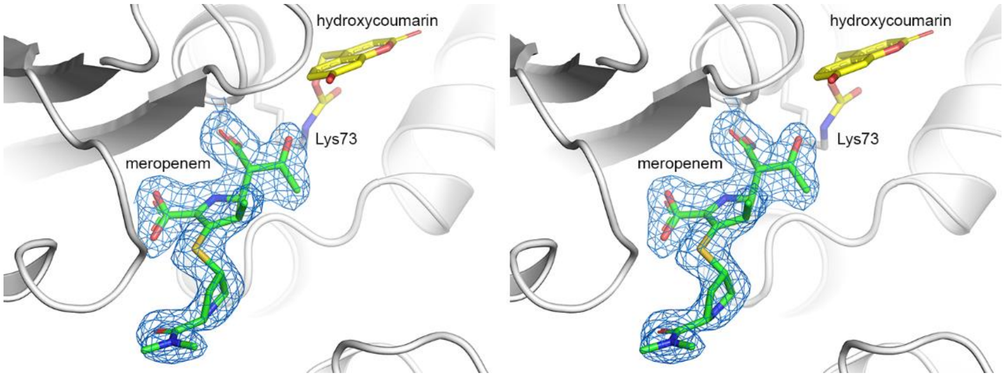


**Supplementary Fig. S7. Meropenem soaked into HC73 crystals.** Stereoview of the $HC73_B$ active site showing that the HC photocage (yellow sticks) remains essentially intact and there is strong electron density (blue mesh, 1σ) for an acylated meropenem (green sticks)

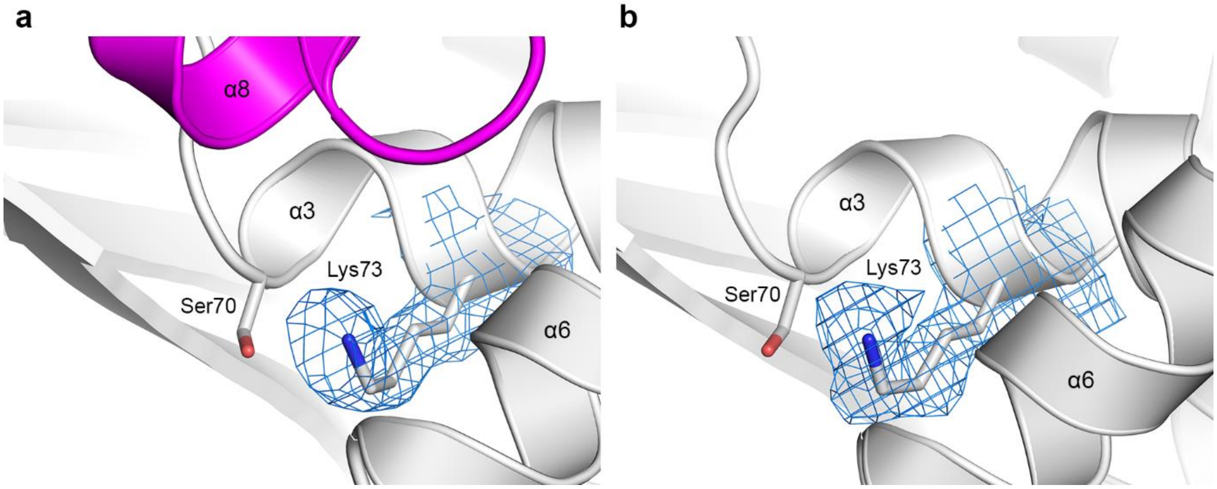


**Supplementary Fig. 8. UV decaging control structures. (a)** The $HC73_A$ active site showing $2F_o$-$F_c$ electron density for Lys73 following HC release. **(b)** The $HC73_B$ active site showing $2F_o$-$F_c$ electron density for Lys73 following HC release. In both structures, the electron density at Lys73 appears marginally larger than expected for an unmodified lysine side chain. This additional density may indicate incomplete photocage removal, resulting in residual occupancy of the hydroxycoumarin linker, or alternatively reflect partial formation of a modified lysine species, such as carboxylation of the Nζ atom. Although the density does not allow unambiguous assignment, the observation suggests coexistence of decaged Lys73 and a fraction of carboxylated Lys73, consistent with re-establishment of the catalytically relevant lysine carboxylation state following photocage release

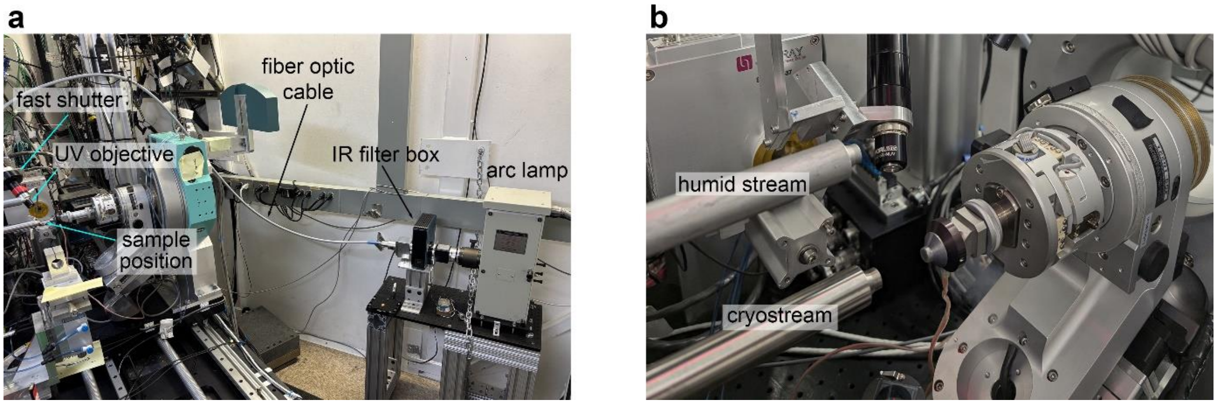


**Supplementary Fig. S9. CAGE-TRX experimental setup at BL7-1. (a)** Output from a 1000 W Newport arc lamp is directed through an IR filter, fiber optic cable, fast shutter, and UV objective onto the sample position. **(b)** A mechanized cryo-trapping system developed in-house allows for rapid switching between a humid air stream and a cryo stream at 100 K.

**Supplementary Notes**

**Supplementary Note 1. Details of the class A and class D β-lactamase active sites.**

The class A and class D β-lactamases are serine-dependent hydrolases responsible for high-level bacterial resistance to β-lactam antibiotics (penicillins, cephalosporins and carbapenems). Although the enzymes from these two classes share a similar 3-dimensional structure (Supplementary Figures 2d and 2e), they have limited amino acid identity with the exception of two highly-conserved sequence motifs directly related to their hydrolase activity(refs). Both enzymes have two structural domains: Domain 1 is composed of a mostly antiparallel β-sheet with two flanking helices, and Domain 2 is a globular α-helical domain with two extended surface loops (the P-loop and the Ω-loop). The active site is located in a cleft between the two domains, with two important catalytic residues, Ser70 and Lys73, located at the N-terminus of an α-helix in the first conserved serine β-lactamase sequence motif SxxK. In both classes of enzyme, Lys73 is thought to play a role in activating the catalytic serine by abstracting a proton[4], thus making it an ideal choice for UAA mutagenesis and photocaging. In the apo form of the enzymes, the lysine is typically hydrogen bonded to the serine side chain, poised to carry out this function. In class D enzymes, Lys73 is post-translationally modified by carboxylation (Supplementary Figure 2c), a modification essential for the deacylation step[5]. To prevent decarboxylation, the lysine is sequestered in an internal the catalytic lysine pocket (CLP) formed by residues from helices α3, α6, and the Ω-loop. The second lysine chosen for UAA mutagenesis (Lys234 in GES-5 and Lys208 in OXA-48) is adjacent to the catalytic serine, located in the highly-conserved KTG sequence motif on strand β5 [6]. This lysine bridges two sides of the active site cleft through a hydrogen bonding interaction with a second conserved serine (Ser130 in class A and Ser118 in class D). This interaction also serves to anchor this serine in the correct orientation to act as a proton shuttle during the acylation step[4,7].

**Supplementary Note 2. The structures of wild-type substrate-free OXA-48 (WT) and HC photocaged apo-OXA-48-K73HCK (HC73)**

The wild-type apo-OXA-48 (WT) and the HC photocaged apo-OXA-48-K73HCK (HC73) mutant were crystallized under similar conditions to those in earlier studies[8,9]. Despite this, the crystals obtained here exhibited different space groups and unit cell parameters (Supplementary Table 2), both relative to those previously reported and between the WT and HC73. This variability is consistent with earlier observations that OXA-48 displays a strong propensity to crystallize in multiple lattice forms, with distinct unit cell parameters arising even under ostensibly identical conditions[9].

The WT and HC73 structures comprised four monomers (A - D) in their respective asymmetric units, arranged as a dimer of dimers (A/B and C/D). Dimerization of OXA-48 has been previously demonstrated, and its high propensity to form dimers is attributed to strong complementarity at the interface[10]. The quaternary assemblies of WT and HC73 are shown in Supplementary Figs. S4a and S4b. While the individual dimers are highly similar between the two structures, the relative packing of the two dimers differs, giving rise to distinct crystal lattices. Formation of each dimer (A/B or C/D)( Supplementary Fig. S4c) buries approximately 3000 $A^2$ of surface area per monomer corresponding to ~30% of the total solvent-accessible surface area in both structures (Supplementary Fig. S4d). This substantial buried surface area is consistent with a stable dimer interface and likely contributes to the observed robustness of dimer formation across different crystal forms.

Each independent WT monomer comprises 241 residues (Trp25 - Pro265) folded in two distinct domains, a β-sheet domain and an α-helical domain with the active site sandwiched between. The catalytic residues (Ser70 and Lys73) are located at the N-terminal end of helix α3 (Figure 2a). The active site contains several well-ordered water molecules, including one hydrogen bonded to the amide nitrogen atoms of Ser70 and Tyr211 in what is known as the oxyanion hole in the serine β-lactamases[11] (Supplementary Figs S2b and S2c). There was a broad peak in the residual $F_o$-$F_c$ electron density at the end of the Lys70 side chain which was modeled as a carboxylate group covalently attached to the Nζ atom.

The arrangement of each HC73 dimer (A/B or C/D) and their respective dimer interfaces are similar to the WT dimer. However, in contrast to the WT structure, the monomers which comprise the HC73 dimers do not have identical structures. Monomers A and C comprise the same 241 residues (Trp25 - Pro265) as their WT counterparts, but monomers B and D have 15 residues missing from a loop between Asn146 and Ile162. In the intact OXA-48 structure (and in all known class D β-lactamases) this amino acid sequence between helices α7 and α9 is known as the Ω-loop (Fig. 1h) and comprises helix α8 harboring two residues (Trp157 and Leu158) critical to the formation of the CLP and thus to enzyme activity[6,12-14].

Most notably, the position of the HC cage is significantly different between the two monomers. In $HC73_A$, which closely resembles the $WT_A$ structure, the HC photocage occupies the enzyme active site (Fig. 2b), where it is stabilized by hydrophobic interactions with Ile102, Trp105 and Tyr211, and a hydrogen bond between the C7 hydroxyl and Thr213. Occupation of the active site displaces the catalytic serine (Ser70) side chain, forcing it into a rotamer conformation intermediate between the standard *p* and *m* states[15]. Although this conformation is typically disfavored, it is stabilized in this instance by a hydrogen bond with the backbone amide nitrogen of Tyr211. Aside from this localized adjustment, incorporation of the HC photocage does not significantly perturb the rest of the active site or the overall protein structure. Conversely, in the $HC73_B$ the HC photocage is flipped into a different conformation, primarily a rotation of ~90° about the Cε-Nζ bond, and occupies a position which would overlap Trp157 and Leu158 in monomer A (Fig. 2c). A hydrophobic interaction with Val120 and a hydrogen bond from Asn124 to the C2 ketone oxygen anchors the photocage. This results in the observed disorder of 15 residues of the Ω-loop, a structural feature of the class D β-lactamases generally deemed to be stable despite its location on the molecular surface[16]. Moreover, the alternate arrangement of the photocage results in an unoccluded active site and the Ser70 side chain adopts a wild-type *m* rotameric conformation.

**Supplementary Note 3. Stability of the WT and HC73 dimers.**

To quantify and compare local and global differences between the HC73 and WT dimers, 250 ns molecular dynamics (MD) simulations were conducted on the individual monomers from each structure. The models were prepared with Maestro (Schrodinger) using the OPLS3e force field[17], and the pre-defined TIP3P water model[18] was used to build the system. The overall charges of the models were neutralized with Na+ and $Cl^-$ ions, and 0.15 M NaCl was added prior to building the system. The systems were minimized prior to the final 100 ns production step run at 300 K and 1 Atm pressure, using the Nosé–Hoover chain coupling scheme for temperature control and the Martyna–Tuckerman–Klein chain coupling scheme with a coupling constant of 2.0 ps for pressure control[19]. Nonbonded forces were calculated using an r-RESPA integrator. The MD simulations were performed using Desmond[20] in the Schrodinger 2019-2 release, and the trajectories were saved at 25 ps intervals for analysis. Analyses were carried out in Maestro using the Simulation Interactions Diagram and Simulation Event Analysis. Maestro and Desmond were run on the SHERLOCK 3.0 HPC cluster at Stanford University.

From the resultant MD trajectories, root-mean-square deviations (RMSDs) of the protein backbone relative to the initial frame were calculated to give a measure of overall protein stability. In the WT structure, both monomers were stable over the entire 250 ns trajectory, with $WT_B$ (average RMSD ~1.5 Å) exhibiting marginally more dynamic behavior than $WT_A$ (average RMSD ~1.2 Å)(Supplementary Fig. S5a). A difference was also observed for HC73, where $HC73_A$ (average RMSD ~1.4 Å) was less dynamic than $HC73_B$, with the latter showing increases in RMSD over the first 50-75 ns which plateau at ~2.0 Å (Supplementary Fig. S5b).

The position of the HC photocage exhibited distinctly different behavior in the two HC73 monomers. In $HC73_A$, the photocage remained stably positioned throughout the 250 ns simulation (blue trace; Supplementary Fig. S5c). Four key distances between the cage and surrounding protein residues were monitored (Supplementary Fig. S5d), revealing two distinct phases: (i) an initial phase spanning the first 75 ns and (ii) a second phase covering the remainder of the trajectory (Supplementary Fig. S5e). During the first phase, the photocage undergoes a reorientation driven primarily by the immediate flip of the carbamate moiety towards the α5-α6 loop. Additional rotations about the O42-C41 and C41-C4 bonds invert the orientation of the γ-lactone ring relative to the crystal structure and displaces the phenolate ring such that the contact between O7 and Thr213 is lost. At ~80 ns rotation about the O42-C41 bond brings the HC photocage back toward strand β5 where the O2 atom forms stable hydrogen bonding interactions with Arg250 that persist for the duration of the trajectory. Throughout the entire 250 ns simulation, the photocage remains confined within the active site, adjacent to Ser70, consistent with continued steric hindrance of substrate access and prevention of acyl-enzyme intermediate formation (Supplementary Fig. S5f). In contrast, the HC photocage in $HC73_B$ displays substantially greater mobility (orange trace; Supplementary Fig. S5c). The initial position (as seen in the crystal structure) is seemingly stable for the first 5-10 ns but the cage then subsequently undergoes significant displacement, moving an average of ~8 Å from its starting position from ~15 ns onward. Notably, the photocage never enters the active site during the simulation.

To assess residue-specific dynamics resulting from HC photocage incorporation, root-mean-square fluctuations (RMSFs), which quantify the average positional deviation of each residue from its mean location over time, were calculated for the WT and HC73 monomers. The $WT_A$ and $WT_B$ monomers show a similar degree of structural flexibility (Supplementary Fig. S6a), whereas the HC73 monomers exhibit divergent behavior (Supplementary Fig. S6b). As expected, well-ordered secondary structure elements exhibit the lowest fluctuations, while higher flexibility is observed in loops, turns, and at the N- and C-termini. Notably, three large external loops critical for enzymatic function[16], the P-loop, the Ω-loop and the B-loop (Fig. 1h) exhibit elevated mobility.

$HC73_A$ closely resembles $WT_A$ in its RMSF profile, as illustrated in the RMSF difference plot comparing the two monomers (Supplementary Fig. S6c). In $HC73_A$, reduced P-loop mobility is directly linked to the presence of the HC photocage. This loop which forms one side wall of the class D β-lactamase active site, is known to exhibit conformational adaptability both in response to substrate binding[7,21,22] and to structural changes in the neighboring α5-α6 loop induced by the presence of small molecules in the active site[23,24]. A similar effect is observed when the carbamate group of the HC photocage flips toward the α5-α6 loop, exerting steric pressure that pushes the loop away from the catalytic serine (Supplementary Fig. S5f). This movement is coupled to an outward displacement of the P-loop, mediated by hydrophobic interactions between Trp105 and Val120 on the α5-α6 loop[16], creating space in the active site for the HC photocage to test alternate conformations (Supplementary Figs. S5f and S6d). The phenolic ring of the cage maintains contact with the Trp105 side chain and limits P-loop.

In contrast, $WT_A$ lacks substrate interaction with Trp105, resulting in greater P-loop fluctuations (Supplementary Fig. S6c).

The $HC73_B$ monomer displays substantially increased flexibility relative to $WT_B$ (Supplementary Fig. S6e). Although RMSFs could not be calculated for the disordered Ω-loop, all other regions show significantly elevated fluctuations relative to $WT_B$. Loss of the Ω-loop effectively untethers helices α6 and α7 from the rest of the enzyme, rendering both helices unusually dynamic. These motions are coupled to substantial displacements of the HC photocage, which responds to shifts in helix α6 (Supplementary Fig. S6f). In addition, disruption of the contact between the Ω-loop and the B-loop, deemed critical for the CHDL activity of OXA-48[16], leads to a marked increase in B-loop flexibility (Supplementary Fig. S6e).

**Supplementary Note 4. $HC73_B$ meropenem complex.**

In the WT enzyme, acylation is initiated by the abstraction of the proton from the Ser70 hydroxyl by the carboxylated lysine, followed by nucleophilic attack on the β-lactam carbon to form a reversible tetrahedral intermediate[4,25]. The proton is then shuttled via Lys73 and Ser118 to the N4 atom of the tetrahedral intermediate, ultimately cleaving the C-N bond of the β-lactam to give the stable acyl-enzyme intermediate (Supplementary Figure 2a). In $HC73_B$, the caged Lys73 has the equivalent of a carboxylate linking the Nζ of the lysine with the HC group, so in this case can still be the proton acceptor and shuttle it to Ser120. Soaking of meropenem into HC73 crystals under red light conditions, and subsequent structure determination showed strong electron density in $HC73_B$ for an acylated meropenem (Supplementary Figure 7). Conversely, in $HC73_A$, where the HC photocage occluded the active site, no evidence of acylation of Ser70 was discernible.

Although unexpected, the presence of both an active and an inactive form of the enzyme in the same crystal gives us a positive control for subsequent decaging and TRX experiments. The concurrent presence of two different enzyme structures *in crystallo* also raises an important question: did the disruption of the Ω-loop and the binding of the HC photocage in its place to produce an active mutant enzyme occur during protein translation, later in solution, or during crystal formation? The activity assays on HC73 show that all of the expressed and purified HC73 was inactive, and that UV illumination produced activity roughly equal to WT. This would argue against there being two populations in the sample, one inactive ($HC73_A$-like) and the other active ($HC73_B$-like). Given the differences in crystal packing between WT and HC73, it is likely that the disruption of the Ω-loop occurred during crystallization.

The ability of $HC73_B$ to effectively hydrolyze meropenem, while $HC73_A$ could not, suggests that the HC photocage in $HC73_A$ is the primary cause of its inactivity. Although no meropenem was observed bound near the active site of $HC73_A$, it is presumed that, given the abundance of substrate in the medium, meropenem could readily enter the active site once the HC photocage was lysed. Thus $HC73_A$ has the potential to be an ideal starting structure for TRX experiments. The first step in testing this premise was to determine whether HC decaging could take place *in crystallo*. Although UV decaging of *o*-(2-nitrobenzyl)-L-tyrosine introduced into lysozyme by UAA mutagenesis as been demonstrated *in crystallo* [26], decaging of HCK in crystals has not been reported. A crystal of HC73 was irradiated with a UV flash at room temperature and the crystal flash-cooled at 100 K. Analysis of the active sites of the $HC73_A$ and $HC73_B$ monomers showed that in both cases the HC photocage was no longer attached to the Lys73. Residual $F_o$-$F_c$ density near the Nζ atom suggested that the lysine residues were now present in their carboxylated form, consistent with cleavage of the C41-O42 bond (Fig. 1c) and the inherent stability of carboxylated Lys73 under the conditions used for crystallization.
